# Low-dose cytokine immunotherapy of solid cancers enabled by phagocytic-competent protein co-crystals

**DOI:** 10.1101/2023.04.06.534711

**Authors:** Michael H Jones, Nirk E Quispe Calla, Robert Smith, Callum Talbot-Cooper, Simon Rudge, Hiroyuki Kusano, Takayuki Shiomi, Yuichi Ishikawa, Hong Zeng, Jonathan Best

## Abstract

Protein therapeutics are often compromised by sub-optimal biodistribution contributing to poor efficacy and adverse events. Drug delivery mechanisms better able to target protein drugs to the disease site and provide localized, sustained release have the potential to transform therapeutic standards. PODS^®^ crystals (PODS) are natural-mimetic, micron-scale protein co-crystals engineered to incorporate a protein cargo that can be sustainably released under the action of resident proteases. PODS are efficiently taken up by phagocytic cells with the cargo protein subsequently released in a bioactive form. Since blood-circulating phagocytic cells, including monocytes, are actively recruited into diseased and inflamed tissue, such as the tumour microenvironment, we postulated that monocyte/macrophage-mediated PODS delivery could be used as a molecular “Trojan horse” to efficiently deliver therapeutic proteins to target cells. This could improve the pharmacodynamics and pharmacokinetics of protein drug delivery to treat systemic and disseminated diseases. Interleukin-2 (IL-2) is notoriously toxic at the high doses required for therapeutic efficacy. Here, we demonstrate the therapeutic efficacy and tolerability of low doses of PODS containing IL-2 cargo (PODS-IL-2) administered intravenously in a mouse model of melanoma. We further demonstrate the therapeutic benefit of PODS delivering IL-2, interleukin-15 (IL-15) and interferon gamma (IFN-γ) in a mouse model of renal cell carcinoma at two doses. Efficacy was seen in both doses with the higher dose generating rapid and complete rejection of the tumour in some of the mice treated with each cytokine. This study provides proof-of-concept for the utility of intravenously administered PODS to provide a generalised and widely applicable mechanism to effectively deliver protein drugs for the therapy of cancer and potentially other diseases.

## Introduction

The therapeutic potential of immunomodulatory proteins, including cytokines^1^ and checkpoint inhibitors^2^, are undermined by dose-limiting toxicity and a high frequency of adverse events. For example, high-dose interleukin-2 (hIL-2), used to activate cytotoxic CD8+ T-cells (T cells) for the treatment of metastatic renal cell carcinoma (mRCC) and melanoma, suffers from wide-ranging and severe side effects including vascular leak syndrome^3^. This toxicity limits the use of hIL-2 as a front line therapeutic.

Following intravenous (IV) administration, IL-2 has a half-life in circulating blood of around 4 minutes^4^. This short half-life is primarily a result of active filtration by the kidneys. Consequently, hIL-2 is administered either by continuous infusion, subcutaneous or intraperitoneal injection^4^. These administration strategies effectively increase IL-2’s half-life in the blood to several hours. IL-2 then inefficiently diffuses from blood to the tumour microenvironment (TME) to reach and activate target T cells.

As illustrated by IL-2, systemic therapy with immunomodulatory proteins is challenging because the pharmacokinetics of these unstable proteins requires high protein concentrations in the serum to deliver even small therapeutic doses to the target tissue. To address this challenge, a drug delivery mechanism that (1) stabilizes the protein (2) maintains low serum levels (3) actively delivers to and targets the diseased tissue and (4) provides sustained release at the target tissue, can be expected to reduce drug-related toxicity and improve therapeutic efficacy.

PODS^®^ (Polyhedrin Delivery System)^5^ is a natural mimetic technology that incorporates target proteins into nano-to-micron-scale crystal structures to form PODS co-crystals (PODS). Purified PODS are non-brittle, durable, and transparent. PODS are stable at 37°C in sterile solution, and across a range of pH values ranging from pH4 - pH10. Stability is lost outside this pH range and in the presence of proteases, including matrix-metalloproteases^6^. These proteases slowly degrade the PODS matrix, releasing a stream of bioactive cargo proteins over a period of 4 – 8 weeks. There is no burst release.

The therapeutic potential for sustained release of proteins from PODS has been demonstrated in a mouse model of cartilage repair^7^ and a rat model of bone repair^8^ in which PODS were locally administered. However, for therapy of non-localized targets, including disseminated cancers, localized drug administration is not feasible.

Exploiting monocytes and macrophages and other phagocytic immune cells as part of a molecular “Trojan horse” drug delivery strategy has previously been proposed^9,10^. Attracted by chemokines^11^, monocytes differentiate into macrophages and extravasate from blood capillaries to actively infiltrate diseased and inflamed tissues and orchestrate the immune response. Although the molecular Trojan horse concept is well established, efforts to date to reduce this to therapeutic practice have not been successful^10^.

As may be predicted from their physical characteristics^12^ (size, shape, surface charge and elasticity), we previously reported^5^ that PODS are efficiently taken up by phagocytic cells *in-vitro*. We also demonstrated that, following phagocytic uptake, the PODS cargo proteins are sustainably secreted in a bioactive form that is able to modulate the phenotype of adjacent heterogenous cells.

In the present study, we used two mouse models of sub-cutaneous cancer to show that intravenous injection of PODS, which generate almost undetectable levels of IL-2 in serum, are able to significantly reduce the growth of sub-cutaneous melanoma. Furthermore, we demonstrated that PODS carrying IL-15, and interferon gamma (IFN-γ) each exert a similar therapeutic effect in limiting the growth of sub-cutaneous renal cell carcinoma (RCC). In some cases, this resulted in complete rejection and detachment of the subcutaneous cancer.

## METHODS

### PODS Crystal Production

PODS were essentially prepared as described^8,13^. Each cargo protein utilized the H1 tag for incorporation into the PC. Human homologs of each cytokine were used. Protein sequences are shown in Supplementary Data Table 1.

### Cell Culture and Differentiation

Monocyte suspension cells (THP-1, Public Health England Culture Collection) were cultured and differentiated to macrophages as described previously^5^

### Measurement of the release of hIL-2 from PODS-hIL-2, with and without macrophages

THP-1 cells were seeded at a density of 2×10^5^ cells/ml in 96-well plates and differentiated into M0 macrophages. M0 cells were then incubated with 1×10^5^, 2×10^5^, or 4×10^5^ PODS IL-2 per ml for 24 h in complete medium (resulting in 5, 10, or 20 PODS/cell). Additionally, the same number of PODS containing human IL-2 (PODS-IL-2) were added to the wells of a 96-well plate without cells and spun down at 3000 x g for 25 min. Cells were then washed twice with PBS, and fresh complete medium was added. After incubation, the medium was collected at 0, 1, 2, 3 and 6 days and subsequently tested by IL-2 ELISA (R&D Systems) according to the manufacturer’s protocol.

#### Cancer cell culture

The B16-F10 melanoma cell line was obtained from ATCC (CRL-6475™). Cells were expanded in Dulbecco’s Modified Eagle’s Medium (Corning^®^) containing 10% fetal bovine serum. The RENCA cell line was obtained from ATCC (CRL-2947) and cultured as above.

#### Generation of tumours

All animal procedures were conducted in compliance with the Stanford Administrative Panel for Laboratory Animal Care (APLAC). For the melanoma model, 6- to 8-week-old female C57BL/6 mice were purchased from Jackson Laboratories. 1×10^5^ B16-F10 cells in 100 μL of PBS were injected subcutaneously into the shaved lateral flank of mice anesthetized by inhalation of isoflurane. For the RCC model, 6-to 8-week-old female BALB/C mice (Jackson laboratories) were used. 2.5×10^5^ RENCA cells were injected subcutaneously as above.

### PODS administration in melanoma model

Fifteen C57BL/6 mice (Jackson Labs) were divided into three groups of five mice in which each mouse the control group either had no injection or received and injection of 4×10^7^ PODS containing enhanced green fluorescent protein (PODS-GFP) in 200μL of PBS. The treatment group received an injection of 4×10^7^ PODS-IL-2 in 200μL of PBS. Intravenous injections were given on days 5, 7, 13, and 19 days after tumour inoculation (total 16×10^7^ PODS per mouse).

### PODS administration in the renal cell carcinoma model

Two separate dosing experiments were conducted. In each experiment Twenty BALB/C mice (Jackson Labs) were divided into four groups of five mice. In the first experiment, each mouse in the treatment groups received an injection of 4×10^7^ PODS containing either PODS-IL-2, PODS-IL-15, or PODS-IFN-γ in 200μL of PBS (total 16×10^7^ PODS per mouse). The remaining group of five mice were untreated controls.

In the second experiment each mouse in the treatment groups received an injection of 1×10^7^ PODS containing either PODS-IL-2, PODS-IL-15, or PODS-IFN-γ in 200μL of PBS. Human homologs of each cytokine were used. Intravenous injections were given on days 7, 10, 16, and 23 days after tumour inoculation (total of 4×10^7^ PODS per mouse). The remaining group of five mice were untreated controls.

### Tumour measurement

Blinded assessment of ulceration around the tumour site were made, and measurements of tumour length and width were performed using external callipers. Tumour volume (mm^3^) was calculated by the modified ellipsoidal formula: V = ½ (Length × Width^2^). Animals were sacrificed if ulceration of the tumour site was observed.

### ELISA measurements of IL-2 in serum

30 μL blood samples were collected at 24 hours after the final dose from C57BL/6 mice receiving 4×10^7^ PODS-IL-2 treatment carrying the B16-F10 melanoma tumours and at 4, 24 and 48 hours intervals from 3 BALB/C mice carrying RCC tumours receiving 1×10^7^ PODS-IL-2 treatment following the final dose and tested by IL-2 ELISA (R&D Systems) according to the manufacturer’s protocol.

#### Immunohistochemical methods

After deparaffinization and rehydration, tumour sections were incubated in a 3% H_2_O_2_ solution for 10 minutes to reduce non-specific background staining. Tissue samples were heated in a 10mM Tris/ 1mM EDTA buffer pH9. Slides were incubated for 30 minutes at room temperature with a primary antibody for CD8a (Invitrogen 4SM16) diluted x100. The primary antibody was visualized using Simple Mouse Stain MAX-PO (Rat) (Nichirei Biosciences, Tokyo, Japan) for 30 minutes, followed by staining with 2% Giemsa solution (Muto Pure Chemicals 15003, Tokyo, Japan).

#### Pathological evaluation

During the pathological evaluation, the staining rate was determined for 10 randomly selected areas at 40x magnification (high power fields, HPF) in each tumour by an experienced senior pathologist.

#### Statistics

Survival rates were depicted using a Kaplan-Meier curve. Tumour volume measurements were plotted and analysed using Prism 9.4.1 (GraphPad Software LLC). Two-way ANOVAs were performed comparing the different treatment groups with respect to time. For the melanoma study, individual data points at day 17 were also isolated for the PODS-GFP and PODS-IL-2 groups and compared using an unpaired student’s *t* test. Similarly, in the RCC study, the individual data points and groups at day 22 were isolated and compared using a one-way ANOVA with *post-hoc* Dunnett’s t-test conducted to determine any differences with the control group.

The presence of CD8+ cells from 10 HPFs was observed for each animal in the melanoma study. The individual numbers were plotted for the GFP and IL-2 groups and analysed using Prism 9.4.1 (GraphPad Software, LLC). A two-tailed Mann-Whitney U-test was performed comparing the control and treatment groups.

## RESULTS

### In vitro IL-2 levels

PODS provide sustained release of cargo proteins. The amount of free, soluble protein that accumulates depends on the balance of release and degradation rates. To provide a basic framework for dosing in the in-vivo assays, we measured in-vitro accumulation from PODS containing human IL-2 (IL-2) which were incubated in serum in the presence or absence of THP-1 cell-derived macrophages. Macrophages were allowed 24 hours to take up the PODS. The amount of IL-2 that accumulated in the culture media was measured using an ELISA assay at 0, 1, 2, 3 and 6 days after addition of 10% serum, which provided a source of proteases to break down the PODS matrix liberating the IL-2 cargo. The amount of IL-2 measured was dependent on the number of PODS added (figure 1). The amount that accumulated in the media peaked by day 1 and remained relatively constant until day 6. There was a direct relationship between the number of PODS and the amount of accumulated IL-2 detected by the ELISA assay. As had been observed in our previous study^5^, where human IL-6 release was measured from phagocytosed PODS, ingestion by macrophages significantly reduces the amount of free cargo protein accumulating in the media. In the absence of macrophages, (measured only on day 6) the amount of IL-2 that accumulated was about 10-fold higher.

**Figure 1.**
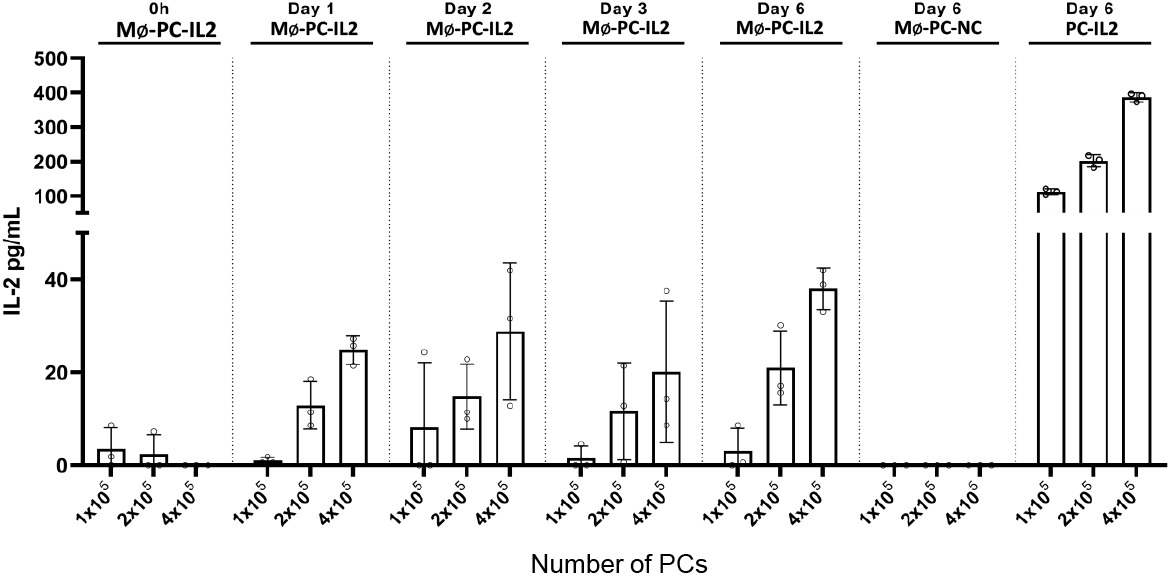
ELISA measurements of accumulated IL-2 released from PODS incubated in the presence or absence of THP-1 macrophages. IL-2 was allowed to accumulate for the number of days indicated in the media before assay. Mø = Macrophages present. NC = no cargo protein in the PODS.

### Determining dosing

To reduce drug toxicity, we wished to limit serum concentrations of IL-2 significantly below levels achieved during treatment of patients with melanoma and mRCC. High dose IL-2 therapy uses 600,000 international units (IU) per kg^14^. 1 mg of IL-2 contains approximately 16 million IU making 600,000 IU equivalent to 37.5 μg/kg. In the clinic, bolus injected patients receive 3 injections per day for 5 days, or as long as they can tolerate. Since residual IL-2 will be available on the second and subsequent injections, peak load is likely to be higher than 37.5 μg/kg.

Based on the above *in-vitro* release data (figure 1), we determined the dosing regimen factoring in mouse weight and published estimates of blood volume and monocyte numbers^15^. For the melanoma study, the mice were tail vein injected with 4×10^7^ PODS per injection. Each mouse was scheduled to receive four injections (a total of 16×10^7^ PODS).

Based on the levels of accumulation seen in-vitro following ingestion of PODS by macrophages, the maximal amount of hIL-2 released would be 16 ng. For a typical 20 g mouse with 2 ml blood volume, this total has the potential to generate a maximum bio-available load of 0.8 ug/kg (about 50-fold lower than the typical hIL-2 concentrations of 37.5 μg/kg) or 8 ng per ml of serum with a maximal burden of 25-75 crystals per circulating monocyte.

In a separate experiment, groups of RCC mice also received a reduced dose of 1×10^7^ PODS per injection and a total of four doses (a total of 4×10^7^ PODS). This total has the maximum potential to generate 0.2 ug/kg or 2 ng per ml of serum with a maximal burden of 6-19 PODS/circulating monocyte.

### Melanoma response to PODS-hIL-2

The melanoma mice were divided into three groups, each of five mice. The first control group received no treatment, the second control group received PODS containing green fluorescent protein (GFP) and the final group received PODS-IL-2.

All 15 mice appeared to be generally healthy with no signs of cachexia throughout the study. The tumours in the mice in the two control groups (GFP and no injection) grew faster than tumours in the PODS-IL-2 treatment group (figure 2A). Consistent with low levels of immunogenicity and bioactivity previously reported for PODS lacking bioactive cargo proteins^16,17^, the mice receiving PODS-GFP did not have any apparent differences in overall health compared with the no-treatment group.

**Figure 2.**
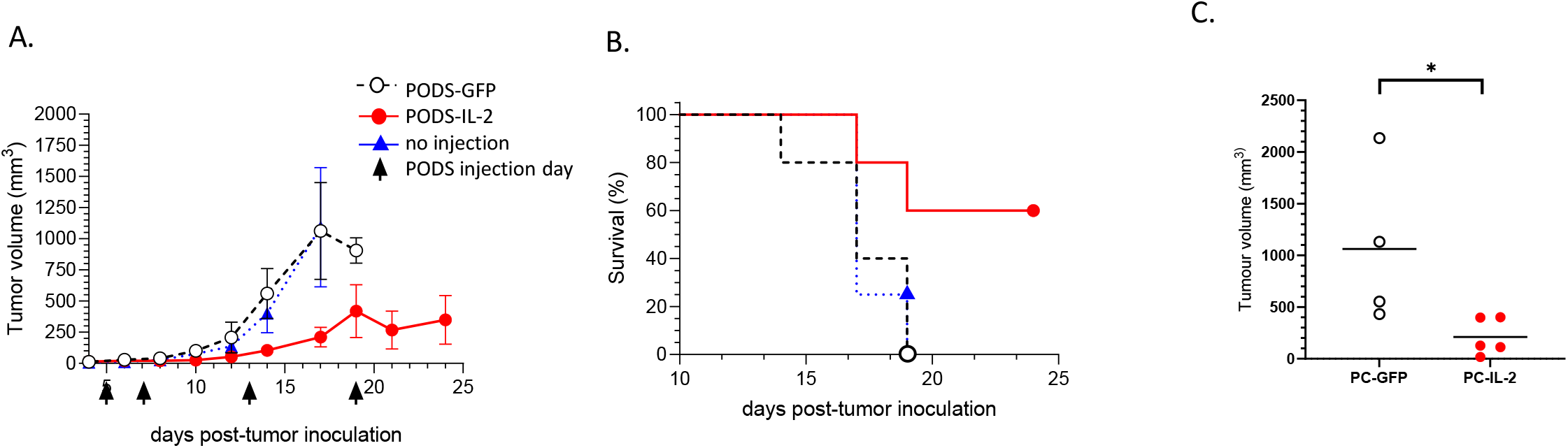
Survival and tumour size in a subcutaneous melanoma mouse model in groups of mice treated with PODS-IL-2, PODS-GFP and untreated (no injection). (A) Mean survival plot over time for each group. B) Mean tumour size (± Standard error of the mean) over time per group. C) Comparison of tumour size between groups containing PODS eGFP (n=4) vs PODS IL-2 (n=5) at day 17, line represents mean value; *= p<0.05.

The first control mouse was sacrificed 14 days after melanoma cell inoculation. Survival of all the mice is plotted in Figure 2B. By day 19, all ten control mice (no injection and PODS-GFP injection) had been sacrificed. In contrast, the first mouse in the treatment group wasn’t sacrificed until day 17 and 3 of the treatment group mice survived to day 25 when the study was terminated. In two of the surviving mice, the tumour volume was stable and remained below 200 mm^3^. A two-way ANOVA comparing tumour size in the PODS-GFP and “no injection” group (Figure 2A) was conducted using a mixed-effect model with matching time point values. There was an effect of time (F = 14.31, p=0.003), but no effect of treatment (F= 0.1027, p=0.76). Thus, the presence of the PODS containing the inactive GFP cargo protein did not influence the tumour growth compared to the no injection group. In contrast, a two-way ANOVA comparing the PODS-GFP and PODS-IL-2 treatment on melanoma tumour volume (Figure 2A) confirmed that there was an effect of time (F= 6.43, p=0.005) and an effect of treatment (F=9.264, p=0.016). When combined with observations of the longitudinal data (Figure 2A and B), this demonstrated that the PODS-IL-2 was slowing the growth of the tumour and prolonging the survival of these mice. Due to the reduction in the numbers of mice in the control group towards the end of the study, a meaningful *post-hoc* analysis comparing individual time points could not be conducted. However, to determine if the reduction in tumour volume with PODS-IL-2 treatment showed a statistically significant effect compared to the PODS-GFP group at later time points, a paired t-test was conducted on the data points at day 17 (Figure 2C). This represented the latest time point with the sufficient control mice available for useful statistical comparison. The unpaired, two-tailed test revealed a significant difference between the two groups (t=2.41, p=0.046). The analysis supports the conclusion that administration of PODS-IL-2 reduces tumour growth.

### Renal cell carcinoma response to IL-2, IL-15 and IFN-γ

In addition to treating melanoma, high dose IL-2 has regulatory approval for the treatment of metastatic renal cell carcinoma (mRCC). Therefore, using RENCA cells injected subcutaneously, we evaluated the utility of PODS-IL-2 to reduce tumour growth in a mouse model of RCC. In addition, we explored the efficacy of the PODS-IL-15 and PODS-IFN-γ. Both of these cytokines have been reported to be effective against a range of cancers^17,18^. Human IFN-γ is not cross reactive with the external domain of the mouse IFN-γ receptor^18^. However, it does bind and activate the internal receptor. Since it has previously been reported that phagocytosed crystals release proteins intracellularly^19^, it was our expectation that human IFN-γ delivered using PODS would also be effective in the mouse study.

The design of the RCC study was similar to the melanoma study, but the PODS-GFP control was not used and the strain of mice used was BALB/C. Initially, we used the same dose of 40 million PODS. However, after the second PODS treatment dose was administered, it was apparent that the all of the treated mice (but not the control mice) were suffering from side effects of treatment and the study was terminated early. However, before the experiment was terminated, one of the five mice in each treatment group had completely rejected their tumour which had entirely detached leaving a visible scar (figure 3).

**Figure 3.**
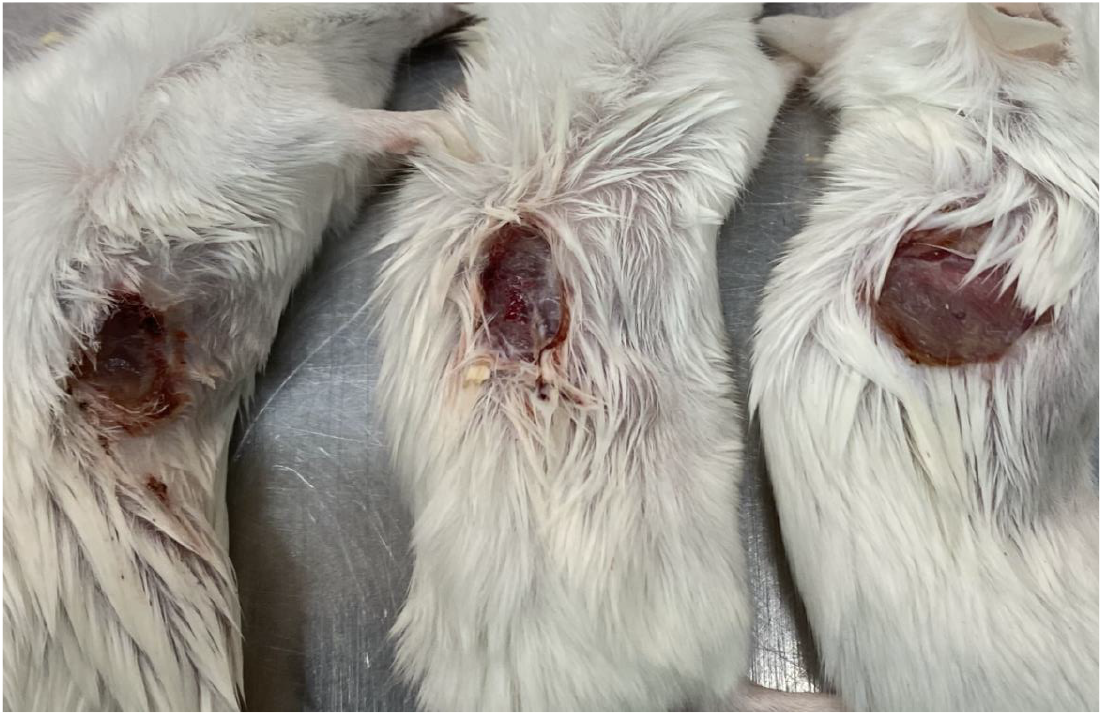
The three mice (one from each treatment group) in which the RENCA tumour detached following two rounds of PODS treatment with 40 million PODS per injection.

Subsequently, the RCC study was conducted using a lower dose of PODS. For each treated mouse, a dose of 1×10^7^ PODS containing a single cytokine cargo was administered a total of four times on days 5, 8, 14, and 20 days after RENCA cell inoculation. Each mouse received a total of 4×10^7^ PODS, 25% of the dose used in the melanoma study and initial RCC study.

Even with the lower drug dose, mice in all three treatment groups showed slower growth (figure 4A) and had longer survival (figure 4B) than those in the untreated group. Two-way ANOVAs were performed comparing tumour size in the different treatment groups with the control group (Figure 4A). Due to the absence of data points for the control group at day 26, this time point was excluded from the analysis. A two-way ANOVA comparing the control group with PODS-IL-2 revealed that there was an effect of time (F=55.62, p<0.0001) and treatment (F=5.63, p<0.05). Equally, there was an effect of treatment when compared with PODS-IL-15 (F=6.56, p=0.014). However, IFN-γ did not show a significant effect with treatment (F=4,79, p=0.06) using this method. As with the melanoma study, the reduction in the numbers of mice in the control group did not allow a meaningful *post-hoc* analysis. Therefore, to determine if the reduction in tumour volume with the treatment groups showed a statistically significant effect compared to the control group, a one-way ANOVA was conducted on the data points at day 22 (Figure 4C). This represented the latest time point with the sufficient mice for useful statistical analysis. The one-way ANOVA revealed a significant effect of treatment (F= 5.68, p= 0.009), with post-hoc analysis compared to the control being significant for all treatment groups (PODS-IL-2, p=0.008; PODS-IL-15, p=0.018; PODS-IFNγ, p=0.024). There was no significant difference when the treatment groups were compared with each other. Although an isolated time point, this result supports the observation that administration of each cytokine results in a reduction of tumour growth.

**Figure 4.**
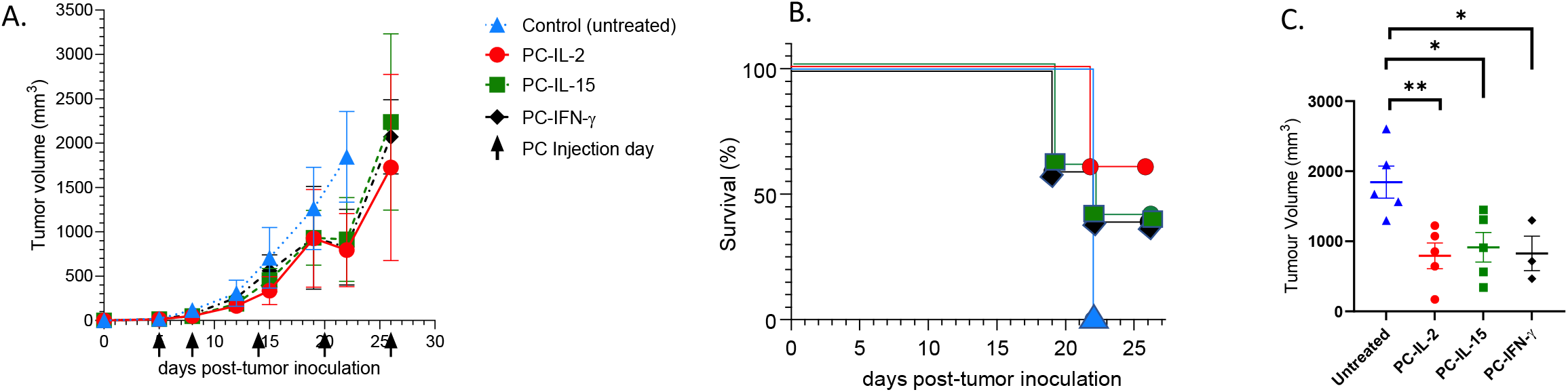
Survival and tumour size in a subcutaneous RENCA mouse model in groups of mice treated with PODS-IL-2, PODS-IL-15, PODS-IFN-γ and untreated (no injection). A) Mean survival plot over time for each group. B) Mean tumour size (± Standard error of the mean) over time per group. C) Comparison of tumour size between the untreated group and groups containing PODS-IL-2 (n=5), PODS-IL-15 (n=5) and PODS-IFN-γ (n=3) at day 22, line represents mean value ± SEM; *= p<0.05, **= p<0.01.

### Activation of CD8+ T cells

The immunotherapeutic effects of IL-2 on cancer are mediated by activation of cytotoxic CD8+ T cells^20^. To evaluate the activity of IL-2 delivered by PODS to the tumour, we examined histological sections from each of the melanomas resected from the mice given PODS-IL-2 or PODS-GFP. Tumour sections were stained with an antibody against CD8 to reveal CD8+ T cells. 30 high powered fields (HPFs) from each tumour were scored blind under the microscope. The numbers of CD8+ cells for each HPF in their respective treatment groups are plotted in figure 5. For all of the PODS-GFP treated HPFs, overall there were less than 20 CD8+ cells seen per HPF. For the PODS-IL-2 treated HPFs, there were multiple fields, derived from two of the tumours in which larger numbers of CD8+ cells were visible. Although there was no statistically significant difference between the groups, these results are supportive of PODS-IL-2 activation of CD8+ T cells.

**Figure 5.**
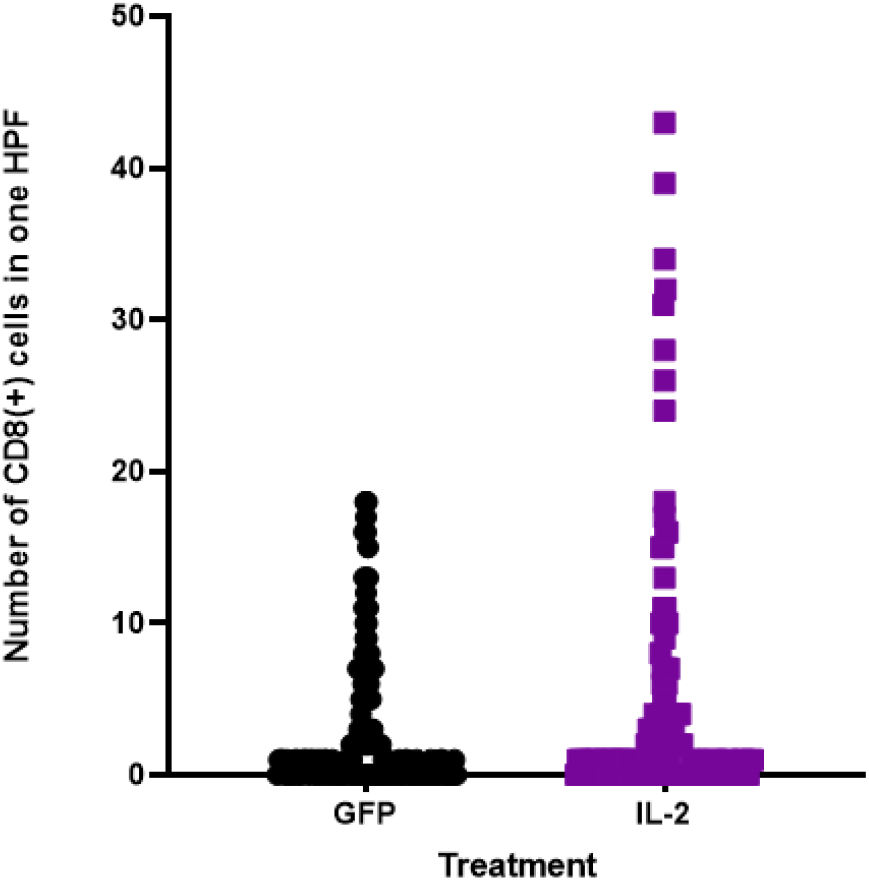
Plot of the number of CD8 positive cells in each high power field (HPF) viewed under the microscope for tumours treated with PODS containing GFP (filled circles) or IL-2 (filled squares).

### Measurement of *in-vivo* IL-2 levels

As well as demonstrating that cytokines delivered by PODS are sufficient to reduce tumour growth, we wished to confirm that serum concentrations were maintained at low levels. Since the conditions in-vivo are very different to in-vitro, it was expected that kidney filtering would maintain low IL-2 serum levels. To measure the amount of IL-2 that actually accumulates in mouse sera, 30 μl blood samples were taken, 24 hours following the final dose, from three of the melanoma study mice that received PODS-IL-2 therapy and three control mice that had received PODS-GFP. For the RCC mice, samples were taken at 4, 24 and 48 hours following the final PODS-IL-2 dose. An ELISA assay was used to measure the amount of free IL-2 that accumulated in the sera (figure 5). Measurements of IL-2 were low, possibly below the limits of assay sensitivity, for GFP and IL-2 melanoma mice. In the IL-2 treated RENCA mice, levels were measured in a range between 9 pg/ml and 17 pg/ml. These values are markedly below the potential maximum levels of 2 ng per ml calculated based on *in-vitro* release rates. This is consistent with constant clearance of released IL-2 in serum by kidney filtration preventing high levels of IL-2 accumulation.

## Discussion

We have shown that PODS, micron scale protein co-crystals, delivering low doses of IL-2 to the TME are able to slow and, in some cases, reverse the growth of melanoma in a mouse model. We also showed a therapeutic effect using IL-2, IL-15 and IFN-γ on RCC in a mouse model.

Addressing toxicity is a high priority for drug developers. Many second-generation IL-2 drugs offering an improved therapeutic window are in development. These employ a variety of mechanisms to provide increased stability, improved specificity, modulating receptor binding characteristics, and tumour targeting^23 24^. Some of these novel delivery modalities are in clinical trials. The delivery system we have described here was designed to exploit active delivery by monocular phagocytic cells to effectively deliver protein drugs to cancer and offers a radically different approach.

A large proportion of the PODS delivered by intravenous injection will lodge directly in other tissues or be filtered, for example by the Kupffer cells of the liver. These off-target PODS will also secrete cargo and, if secretion rates are sufficiently high, cargo proteins secreted from PODS located away from the tumour may also reach the tumour. It is not clear to what extent the therapeutic effect we observed is mediated by the release of IL-2 from PODS that have lodged in the tumour versus IL-2 released by PODS elsewhere in the body. However, given the very low serum IL-2 levels measured, it is unlikely that free IL-2 transported to the tumour, rather than released within the tumour from PODS, was primarily responsible for observed efficacy.

We have not directly demonstrated here that MPs are responsible for delivery to the cancer. However, our in-vitro analysis of PODS uptake by monocytes and macrophages combined with the in-vivo behaviour of MPs makes this a very likely mechanism. Amongst professional phagocytic cells, monocytes (and the macrophages they differentiate into) are the most likely mediators of PODS delivery to tumours. However, it should be noted that neutrophils, mast cells, and dendritic cells also phagocytose efficiently and infiltrate inflamed tissue and cancers^21^. These cells may also contribute to transport of the PODS to the tumour target.

In the RCC mice treated with 40 million PODS per dose, one of each of the tumours in each treatment group vanished following the second round of dosing. Rather than regression, and consistent with a massive immune response, the appearance of the wound left at the site indicated that the tumour had detached from its sub-cutaneous location.

IL-2 provides its therapeutic effect in cancer by increasing T cell activity within the TME. As well as delivering protein to target heterologous cells adjacent to macrophages within the TME, PODS could be used to directly modulate the phenotype of the macrophage itself. Macrophages exist in a spectrum of phenotypes broadly characterised as ranging between classical M1, pro-inflammatory and alternative M2, regenerative phenotypes^11^. Tumours subvert the behaviour of macrophages towards a pro-tumorigenic M2 phenotype. Although useful, the M1-M2 axis is a simplified view of a complex response. IFN-γ can be both pro- and anti-tumorigenic depending on context, but usually has beneficial effects in cancer patients^22^. Human IFN-γ is only cross reactive with the internal domain of the murine receptor^18^. Previous studies by others^19^ have reported intracellular delivery of crystalline proteins following phagocytic ingestion. Our observation of human IFN-γ efficacy in a mouse model of cancer is consistent with evidence that cargo molecules from PODS are also intracellularly delivered and suggest a therapeutic benefit against cancer can be achieved without binding to the extracellular domain of the IFN-γ receptors.

The sustained release of cargo proteins from PODS allows localized concentrations of the cargo protein to accumulate. Since the TME is largely isolated from the filtering effects of the kidney, it can be expected that protein drugs will accumulate in the TME to higher levels than seen in serum. This is the opposite of what normally occurs following intravenous injection where protein drugs perfuse from high serum levels through to markedly lower levels in the target tissue. Detailed studies to track the fate of PODS and their cargos are currently being undertaken and will be important to build an understanding of the adsorption, distribution, metabolism, excretion and toxicity for PODS and their cargo proteins before progress to the clinic.

The amount of IL-2 measured in sera was very low. Unexpectedly, the serum levels in the RCC mice, which received a lower IL-2 dose, were higher than in the melanoma mice at an equivalent time point. Whilst the tumours may have an impact on serum levels, the observation may also be associated with the different genetic background of the two mouse strains with C57BL/6 mice used for the melanoma study and BALB/C mice used for the RCC study. This higher level correlates with the adverse effects seen in the BALB/C mice treated with the higher dose, which was absent in the C57BL/6 mice. Further studies will be needed to understand the underlying mechanism.

As an alternative to direct injection of PODS, efficiency of therapy could be enhanced by introducing PODS to phagocytic immune cells ex-vivo. After a period of incubation to allow uptake of the PODS, the loaded phagocytic cells would be returned to the patient. Compared to other autologous cell-based therapies, such as CAR-T therapy, such a procedure would require very simple manipulation of the cells and would be expected to reduce the number of free PODS that are filtered by Kupffer cells or lodge in non-target tissue enabling higher proportions of PODS-loaded cells to reach the target tissue. As well as reducing toxicity, this may further increase efficacy.

It will also be important to build a greater understanding of the mechanism(s) by which cargo protein is released from PODS following uptake by phagocytic cells. Although macrophages and monocytes usually degrade ingested particles, it is possible that the large amount of protein material present in PODS may overwhelm this apparatus. PODS may also be released periodically following cell death allowing transient cargo release opportunities prior to uptake of PODS by other phagocytic cells.

In addition to cytokines, many other protein drugs suffer from poor PKPD profiles. For example, checkpoint inhibitors, presently at the vanguard of immuno-oncology, produce adverse effects in about 40% of patients^2^. These can be severe enough to halt treatment or cause fatality. The drug delivery mechanism demonstrated here could be applied to other protein drugs and other diseases where inflammation actively recruits phagocytic immune cells. Protein drugs that could be delivered by PODS include other growth factors, chemokines, antibodies and antibody variants (e.g. nanobodies), viruses and virus-like particles, enzymes (e.g. hyaluronidase), and peptide drugs. Moreover, combination therapies with protein/protein and protein/non-protein drugs, such as chemotherapeutics, as well as in combination with surgery may provide higher levels of therapeutic efficacy.

The mechanism provided by PODS has many key attributes that make it particularly attractive: Manufacture of PODS is simple, scalable and low cost. Sustained release is achieved over several weeks without burst release. Moreover, PODS are very well tolerated in mouse models, and are 100% protein and therefore biodegradable. This combination of attributes generates a unique ability in PODS to significantly skew the biodistribution of protein drugs to reduce toxicity and increase efficacy.

**Figure 6.**
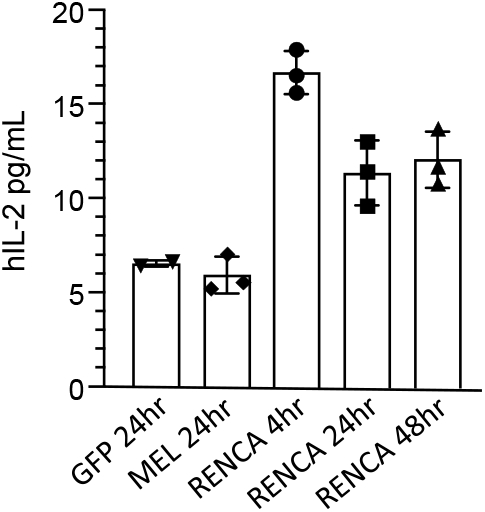
ELISA measurements of serum IL-2 in treated and untreated mice. Blood samples were taken at the time intervals indicated following the final injection of PODS-IL-2 in the melanoma and RENCA mice.

