## Supplementary data for "Low-dose cytokine immunotherapy of solid cancers enabled by phagocytic-competent protein co-crystals"

Supplementary data Table 1

Amino acid sequences. The H1 tag (yellow) and VP3 tags (red) are highlighted

Human IL-2

MADVAGTSNR DFRGREQRLF NSEQYNYNNS KNSRPSTSLY KKAGFAPTSS STKKTQLQLE
HLLLDLQMIL NGINNYKNPK LTRMLTFKFY MPKKATELKH LQCLEEELKP LEEVLNLAQS
KNFHLRPRDL ISNINVIVLE LKGSETTFMC EYADETATIV EFLNRWITFC QSIISTLT

Human IL-15

MADVAGTSNR DFRGREQRLF NSEQYNYNNS KNSRPSTSLY KKAGFNWVNV ISDLKKIEDL
IQSMHIDATL YTESDVHPSC KVTAMKCFLL ELQVISLESG DASIHDTVEN LIILANNSLS
SNGNVTESGC KECEELEEKN IKEFLQSFVH IVQMFINTS

Human IFN-γ

MQDPYVKEAE NLKKYFNAGH SDVADNGTLF LGILKNWKEE SDRKIMQSQI VSFYFKLFKN
FKDDQSIQKS VETIKEDMNV KFFNSNKKKR DDFEKLTNYS VTDLNVQRKA IHELIQVMAE
LSPAAKTGKR KRSQMLFRGR RASQDPAFLY KVVDGYLLAF NSQRRSHTLR LLGPFQYFNF
SETDRGHPLF RLPLKYPSKA IPADELIDNL H
